# Reduction of Nav1.1 in the dorsal striatum preferentially increases hyperthermia-induced generalized seizures compared with the neocortex and nucleus accumbens

**DOI:** 10.64898/2026.05.11.724427

**Authors:** Tetsushi Yamagata, Hiroaki Mizukami, Yurina Hibi, Kazuhiro Yamakawa, Toshimitsu Suzuki

**Affiliations:** Department of Neurodevelopmental Disorder Genetics, Institute of Brain Science, Nagoya City University Graduate School of Medical Sciences, Nagoya, Aichi, 467-8601, Japan; Division of Genetic Therapeutics, Center for Molecular Medicine, Jichi Medical University, Shimotsuke, Tochigi, 329-0498, Japan

**Keywords:** *SCN1A*, Nav1.1, epilepsy, Dravet syndrome, dorsal striatum, nucleus accumbens, febrile seizure

## Abstract

Mutations in *SCN1A*, which encodes the voltage-gated sodium channel Nav1.1 (αI subunit), are the major cause of Dravet syndrome, a severe developmental and epileptic encephalopathy. Although Nav1.1 haploinsufficiency preferentially impairs inhibitory interneuron function, the region-specific contributions of distributed brain circuits to seizure susceptibility, particularly in subcortical structures that have received less attention than the neocortex and hippocampus, remain unclear. Here, we examined the effects of region-specific Nav1.1 deficiency on hyperthermia-induced seizures by selectively deleting *Scn1a* in the neocortex, nucleus accumbens (NAc), and dorsal striatum (caudate–putamen, CPu) of adult mice using an adeno-associated virus-mediated Cre-loxP approach. Contrary to expectations based on prior cortical studies, homozygous *Scn1a* deletion in the neocortex produced only modest effects on seizure generalization. In contrast, homozygous deletion in the NAc and CPu induced generalized seizures to varying degrees. Notably, heterozygous *Scn1a* deletion in the CPu alone was sufficient to trigger generalized seizures, whereas similar manipulations in the neocortex or NAc were not. Seizure threshold temperatures were largely comparable across regions. These findings identify the dorsal striatum as particularly vulnerable to partial Nav1.1 loss and reveal functional heterogeneity within striatal circuits. Our results underscore a previously underappreciated role of striatal inhibitory networks in hyperthermia-induced seizure susceptibility and provide new insights into the circuit mechanisms underlying Dravet syndrome.

## Introduction

Dravet syndrome is a severe developmental and epileptic encephalopathy characterized by the onset of febrile or afebrile clonic and generalized tonic–clonic seizures within the first year of life, followed by the emergence of multiple seizure types, including myoclonic, atypical absence, and focal seizures [Dravet, 2011]. As the disease progresses, patients develop intellectual disability and psychomotor delay, and approximately 10% experience sudden unexpected death in epilepsy (SUDEP). Seizures are frequently refractory to currently available antiepileptic drugs, highlighting the urgent need for mechanism-based therapeutic strategies. Approximately 80% of patients with Dravet syndrome harbor heterozygous loss-of-function mutations in the *SCN1A* gene, which encodes the voltage-gated sodium channel Nav1.1 [Claes *et al*., 2001: Sugawara *et al*., 2002; Lossin, 2009; Escayg and Goldin, 2010; Yamakawa *et al*., 2016]. Haploinsufficiency of Nav1.1 is widely accepted as the primary pathogenic mechanism. Nav1.1 is predominantly expressed in parvalbumin-positive (PV^+^) inhibitory interneurons, and its dysfunction leads to impaired inhibitory neurotransmission and consequent network hyperexcitability [Ogiwara *et al*., 2007; Yamagata *et al*., 2017; Tatsukawa *et al*., 2018; Favero *et al*., 2018]. In addition, *Scn1a* expression has been identified in a subset of cortical projection neurons [Ogiwara *et al*., 2013; Yamagata *et al*., 2023], suggesting that both inhibitory and excitatory neuronal populations may contribute to disease pathophysiology.

Region-specific genetic manipulations have demonstrated that Nav1.1 deficiency in the neocortex or hippocampus markedly increases hyperthermia-induced seizure susceptibility [Stein *et al*., 2019; Jansen *et al*., 2020]. However, whether subcortical structures such as striatum similarly contribute to seizure susceptibility in the context of Nav1.1 deficiency remains largely unexplored, raising the possibility that these regions may play a more prominent role than previously appreciated. This prompted us to investigate whether the striatum contribute to hyperthermia-induced seizure susceptibility under conditions of Nav1.1 deficiency.

The striatum is a key subcortical hub that integrates widespread excitatory inputs from the cortex and modulates motor and cognitive functions through cortico–basal ganglia– thalamic loop [Martel and Galvan, 2022; Shipp, 2016]. Importantly, the striatum is enriched in inhibitory neuronal populations, including PV^+^ fast-spiking interneurons (FSIs), which play a critical role in regulating striatal output and network stability. Emerging evidence suggests that cortico-striatal circuitry is directly involved in seizure generation. Using a mouse model of epilepsy associated with *STXBP1* and *SCN2A* mutations, we previously demonstrated that reduced excitatory synaptic transmission from cortical pyramidal neurons onto striatal PV^+^ FSIs is sufficient to trigger epilepsy [Ogiwara *et al*., 2018; Miyamoto *et al*., 2019]. Consistent with this, pharmacological suppression of cortico-striatal excitatory input onto FSIs has been shown to induce generalized seizures in both rodents and non-human primates [Gittis *et al*., 2011; Aupy *et al*., 2024]. Furthermore, selective suppression of PV^+^ FSIs in the ventral striatum, particularly in the nucleus accumbens (NAc), is sufficient to elicit convulsive seizures [Suzuki *et al*., 2026]. These findings collectively suggest that dysfunction of striatal inhibitory microcircuits, especially those involving PV^+^ FSIs, can critically contribute to seizure initiation and propagation.

Notably, functional heterogeneity along the dorsoventral axis of the striatum has been well documented, with the dorsal striatum primarily involved in motor control, whereas the ventral striatum is implicated in reward processing, motivation, and emotional regulation.

However, whether these subregions differentially contribute to seizure susceptibility, particularly in the context of Nav1.1 deficiency, remains unclear. Given that impaired inhibitory neuron function due to Nav1.1 haploinsufficiency is a central mechanism in Dravet syndrome, we reasoned that Nav1.1 dysfunction in striatal inhibitory circuits may amplify or facilitate the spread of aberrant cortical activity.

Based on these considerations, we first examined whether region-specific Nav1.1 deficiency in the neocortex of adult mice affects hyperthermia-induced seizure susceptibility, as predicted by prior cortical studies. Unexpectedly, neocortical *Scn1a* deletion produced only limited seizure effects compared with previous reports. We therefore generated two independent mouse lines with selective *Scn1a* deletion in either the NAc or the CPu, and systematically compared their respective contributions to hyperthermia-induced seizure susceptibility. We found that homozygous *Scn1a* deficiency in the NAc induced generalized seizures; strikingly, even heterozygous loss of *Scn1a* in the CPu was sufficient to induce generalized seizures, a phenotype not observed following comparable manipulations in the neocortex or NAc. These findings identify the CPu as a critical and previously underappreciated node in modulating temperature-sensitive seizure susceptibility and reveal functional heterogeneity within striatal subregions in the context of *Scn1a* deficiency.

## Results

### Neocortical Nav1.1 deficiency produces limited effects on hyperthermia-induced generalized seizures

To determine whether Nav1.1 deficiency in the neocortex contributes to seizure generation as suggested by prior studies [Jansen *et al*., 2020], we targeted a broader extent of the neocortex, including the motor and somatosensory cortices, which represent regions of high Nav1.1 expression and functional diversity, to more comprehensively assess the contribution of neocortical circuits. We induced region-specific deletion of *Scn1a* in the neocortex of adult mice using AAV-Cre injection and assessed hyperthermia-induced seizure susceptibility 4 weeks later (**Fig. 1**). Selective homozygous *Scn1a* deficiency in the neocortex (*Scn1a*^fl/fl^ with AAV-Cre injection) resulted in a generalized seizures in only one of four mice at 43.1 °C, while two exhibited myoclonic and clonic seizures at 41–42 °C without generalization (**Fig. 1A**). No generalized seizures were observed in heterozygous *Scn1a*^fl/+^ mice with neocortex-targeted deletion. These results were less pronounced than those reported in previous study of cortical *Scn1a* deficiency [Jansen *et al*., 2020], suggesting that the neocortical regions targeted here, including the motor and somatosensory cortices, may not be the primary drivers of hyperthermia-induced seizure generalization.

**Figure 1.**
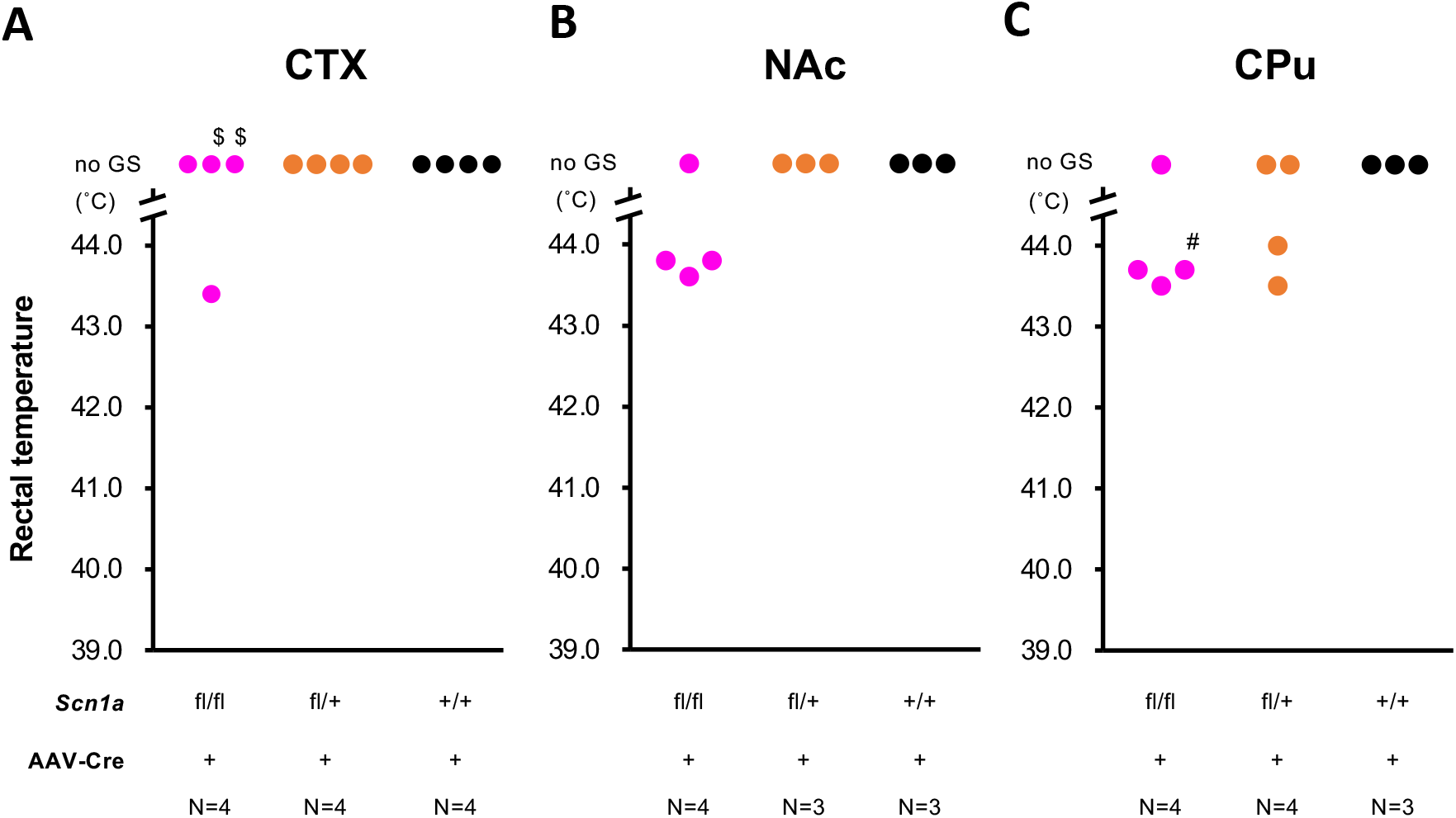
Region-specific Nav1.1 deficiency in the CPu preferentially enhances susceptibility to hyperthermia-induced generalized seizures in 12–13-week-old mice. (**A–C**) Seizure threshold temperatures for generalized seizures following AAV-Cre– mediated deletion of *Scn1a* in the neocortex (CTX) (A), NAc (B), or CPu (C). In contrast to the limited effects observed in the CTX and NAc under heterozygous conditions, generalized seizures were observed in both homozygous and heterozygous *Scn1a*-deficient mice when deletion was targeted to the CPu. Two *Scn1a*^fl/fl^ mice with neocortical deletion did not develop generalized seizures but exhibited frequent myoclonic and clonic seizures at 41–42 °C ($). One *Scn1a*^fl/fl^ mouse died after repeated clonic seizures during the febrile seizure test (#). Dots represent seizure threshold temperatures of individual mice. Sample sizes for each genotype are indicated below the graphs. AAV-Cre, AAV5-Ef1a-mCherry-IRES-Cre; CPu, caudate–putamen; CTX, neocortex; fl, floxed; GS, generalized seizures; NAc, nucleus accumbens.

This unexpected outcome prompted us to investigate whether subcortical regions, specifically striatal subregions such as the CPu and NAc, play a more prominent role.

### Nav1.1 deficiency in the NAc induces generalized seizures under homozygous but not heterozygous conditions

Selective homozygous *Scn1a* deficiency in the NAc (*Scn1a*^fl/fl^ with AAV-Cre injection) induced generalized seizures in three of four mice (mean threshold: 43.7 °C), whereas no generalized seizures were observed in heterozygous (*Scn1a*^fl/+^ with AAV-Cre injection) mice (**Fig. 1B**). These findings indicate that the NAc contributes substantially to hyperthermia-induced seizure susceptibility; however, a complete loss of Nav1.1 function, rather than partial reduction, appears to be required to elicit generalized seizures in this region.

### Heterozygous Nav1.1 deficiency in the CPu is sufficient to induce hyperthermia-triggered generalized seizures

Strikingly, selective homozygous *Scn1a* deficiency in the CPu (*Scn1a*^fl/fl^ with AAV-Cre injection) resulted in a generalized seizure in two of four mice, while one additional mouse died following repeated clonic seizures (mean seizure threshold among affected mice: 43.6 °C) (**Fig. 1C**). Critically, generalized seizures were also observed in heterozygous *Scn1a*^fl/+^ mice with CPu-targeted deletion (2/4; mean threshold: 43.8 °C), a phenotype not observed in heterozygous mice with neocortex- or NAc-targeted deletion, nor in AAV-Cre– injected wild-type mice (*Scn1a*^+/+^) in any region.

Seizure threshold temperatures were comparable across all regions. During the housing period between AAV injection and the hyperthermia test, no spontaneous seizures were noted upon routine observation, and no sudden death occurred in any of the region-specific Nav1.1-deficient mice. Collectively, these findings demonstrated that the CPu is uniquely sensitive to partial Nav1.1 loss and plays a prominent role in hyperthermia-induced seizure susceptibility, identifying the dorsal striatum as a critical node in the epileptic network of Dravet syndrome.

### Verification of AAV targeting to intended brain regions by mCherry fluorescence

To verify the accuracy of region-specific AAV delivery, Cre expression was assessed by mCherry fluorescence. Coronal brain sections were prepared from *Scn1a*^fl/fl^, *Scn1a*^fl/+^, and *Scn1a*^+/+^ mice with AAV-Cre injection (n = 3–4 per group) and examined by fluorescence microscopy. Robust mCherry signals were observed in the neocortex, NAc, and CPu (**Fig. 2**), confirming that AAV was successfully delivered to the intended regions.

**Figure 2.**
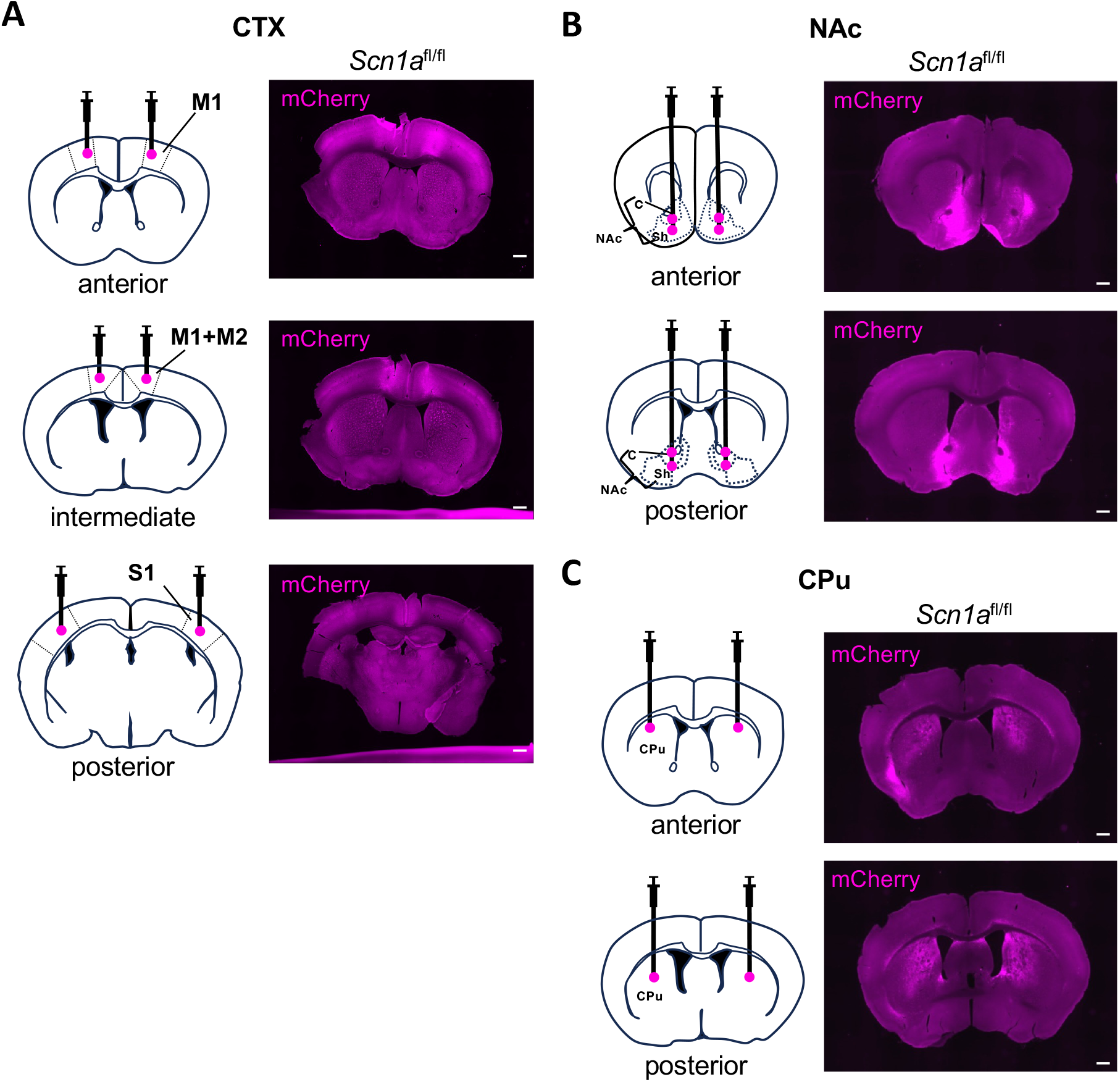
Verification of region-specific AAV targeting by mCherry fluorescence. (**A– C**) Schematic illustration of AAV injection sites in the neocortex (CTX) (A left; magenta circles), NAc (B left) or CPu (C left). Representative coronal sections showing mCherry fluorescence in *Scn1a*^fl/fl^ mouse brain (A–C right), confirming accurate and region-specific AAV delivery. Scale bars: 500 µm. c, nucleus accumbens core; CPu, caudate-putamen; M1, primary motor cortex; M2, secondary motor cortex; NAc, nucleus accumbens; S1, primary somatosensory cortex; sh, nucleus accumbens shell.

## Discussion

In the present study, we investigated the contribution of region-specific Nav1.1 deficiency to hyperthermia-induced seizure susceptibility in adult mice. We initially targeted the neocortex, motivated by prior studies demonstrating that cortical or hippocampal Nav1.1 deletion markedly increases seizure susceptibility [Stein *et al*., 2019; Jansen *et al*., 2020]. However, neocortical Nav1.1 deletion under our experimental conditions produced only modest effects, prompting us to extend our investigation to subcortical striatal regions. Notably, we found that Nav1.1 deficiency in the CPu, even at the heterozygous level, was sufficient to induce hyperthermia-triggered generalized seizures, revealing a prominent and previously underappreciated role for the dorsal striatum in modulating seizure susceptibility.

Region-specific Nav1.1 deletion in the neocortex had limited effects on hyperthermia-induced seizure susceptibility. Generalized seizures were observed in only a subset of homozygous mice and were absent in heterozygous animals. Moreover, seizure threshold temperatures were higher than those reported in previous studies of cortical or hippocampal Nav1.1 deficiency and systemic haploinsufficiency [Stein *et al*., 2019; Jansen *et al*., 2020; Yamagata *et al*., 2020]. This discrepancy may reflect differences in the targeted cortical regions, timing of AAV administration, or observation period. Given the functional heterogeneity of the neocortex, specific subregions may contribute more critically to febrile seizure susceptibility than those targeted here. Therefore, the contribution of cortical Nav1.1 deficiency cannot be fully resolved from the present data.

In the NAc, homozygous but not heterozygous Nav1.1 deletion induced generalized seizures, indicating lower sensitivity to partial Nav1.1 loss compared with the CPu. The ventral striatum is extensively interconnected with cortical and thalamic regions and is involved in limbic and motivational processing. Notably, our previous study showed that reduced PV^+^ FSI activity in the CPu produced only mild epileptiform discharges, whereas similar manipulation in the NAc was sufficient to induce convulsive seizures [Suzuki *et al*., 2026]. At first glance, this appears inconsistent with the present finding that heterozygous Nav1.1 deletion preferentially promotes generalized seizures in the CPu. However, this discrepancy likely reflects differences in the mode and extent of circuit perturbation. Acute suppression of FSI activity may more strongly disrupt ventral striatal circuits, whereas partial Nav1.1 loss may differentially alter network dynamics, predisposing the CPu to mild inhibitory deficits. Regional differences in afferent connectivity, circuit architecture, and compensatory mechanisms may further contribute to these effects. Thus, although the NAc plays a critical role in seizure generation, a greater degree of inhibitory dysfunction may be required to elicit generalized seizures under Nav1.1 haploinsufficiency.

A key finding is that both homozygous and heterozygous Nav1.1 deletion in the CPu resulted in generalized seizures at comparable threshold temperatures. Although the limited sample size precludes firm statistical conclusions, this observation suggests that even partial impairment of Nav1.1-dependent inhibitory function in the CPu may facilitate seizure generalization. The CPu is a central node within the cortico–basal ganglia–thalamic loop, integrating widespread cortical excitatory inputs. Disruption of inhibitory control in this circuit, through reduced excitatory drive onto FSIs or direct suppression of FSI activity, has been shown to produce a graded progression from mild seizures to convulsive seizures [Miyamoto *et al*., 2019]. Together, these findings suggest that diminished excitatory input to CPu FSIs and the resulting disinhibition of striatal output may promote seizure propagation across large-scale networks.

None of the region-specific Nav1.1-deficient mice exhibited spontaneous seizures or sudden death, in contrast to the severe phenotypes observed in the systemic Nav1.1-deficient Dravet syndrome model. This finding suggests that Nav1.1 dysfunction confined to a single brain region is insufficient to recapitulate the full disease phenotype and supports the view that Dravet syndrome arises from distributed network dysfunction involving multiple brain regions and developmental factors. Our results further indicate that the cortico–striatal circuits, rather than isolated cortical dysfunction, may play a key role in shaping seizure expression under hyperthermic conditions.

Several limitations should be noted. First, variability in AAV infection efficiency and spread may have introduced heterogeneity in the extent of Nav1.1 deletion. Second, seizure susceptibility was assessed only using a hyperthermia-induced paradigm, without long-term monitoring for spontaneous seizures. Third, we did not directly evaluate the functional impact of Nav1.1 loss at the cellular or circuit level. Future studies combining in vivo and ex vivo electrophysiological approaches will be necessary to clarify how Nav1.1 deficiency alters neuronal excitability and synaptic integration within cortico–striatal networks. In addition, the contribution of other inhibitory structures warrants investigation. For example, the reticular thalamic nucleus (RTN), composed almost entirely of PV^+^ GABAergic neurons with high Nav1.1 expression, plays a central role in thalamocortical synchronization and may critically influence susceptibility to hyperthermia-induced seizures.

In conclusion, region-specific Nav1.1 deficiency in the adult striatum, particularly in the CPu, preferentially affects seizure generalization rather than markedly lowering seizure threshold temperature. Unexpectedly, neocortical Nav1.1 deficiency had only limited effects on the induction of generalized seizures in our AAV-mediated, region-specific deletion approach, underscoring the importance of examining subcortical circuits in addition to cortical regions. These findings identify the dorsal striatum as an important and previously underappreciated hub within the cortico–basal ganglia–thalamic loop, where disruption of Nav1.1-dependent inhibitory microcircuits, particularly in PV^+^ fast-spiking interneurons, may facilitate seizure propagation across distributed brain networks, thereby advancing our understanding of the pathophysiology of Dravet syndrome. Further studies are needed to define the underlying mechanisms and circuit interactions, but the present results provide a framework for understanding how subcortical regions modulate seizure susceptibility in Nav1.1-related epilepsies.

## Methods

### Mice

All mice were maintained under a 12 h light/dark cycle with ad libitum access to food and water. *Scn1a* floxed (*Scn1a*^fl/+^) mice [Ogiwara *et al*., 2013], in which exon 7 is flanked by loxP sites, were backcrossed onto a C57BL/6J background for more than 10 generations. Heterozygous *Scn1a*^fl/+^ mice were intercrossed to generate wild-type (*Scn1a*^+/+^), heterozygous (*Scn1a*^fl/+^), and homozygous (*Scn1a*^fl/fl^) littermates.

### AAV vector

The pAAV-EF1a-mCherry-IRES-Cre plasmid was obtained from Dr. Karl Deisseroth (Addgene plasmid # 55632) [Fenno *et al*., 2014]. AAV5 packaging, purification, and titration were performed at the Division of Genetic Therapeutics, Jichi Medical University.

### Stereotaxic surgery

AAV injections were performed at 8–9 weeks of age as previously described [Suzuki *et al*., 2024]. Mice were anesthetized with isoflurane (1.0–2.5%) and placed in a stereotaxic apparatus (Stoelting). AAV5-EF1a-mCherry-IRES-Cre (7.3 × 10^12^ viral genomes/ml) was bilaterally injected using a microinjector (Nanoliter 2020; Injector; World Precision Instruments) equipped with a pulled glass capillary at 100 nl/min. Coordinates were based on a mouse brain atlas [Paxinos and Franklin, 2001]. Injection sites were as follows: Neocortex: anteroposterior (AP) +0.98 mm, mediolateral (ML) ±1.20 mm, dorsoventral (DV) -1.63 mm; AP +0.02, ML ±0.80, DV -1.40; AP -1.58, ML ±2.75, DV -1.50 (200 nl/site); NAc: AP +1.60, ML ±1.00, DV -5.00 and -4.50; AP +0.80, ML ±1.00, DV -5.20 and -4.50 (250 nl/site); CPu: AP +0.70, ML ±2.00, DV -3.00; AP 0.00, ML ±2.00, DV -2.80 (500 nl/site).

### Febrile seizure (FS) test

FS tests were conducted in male and female mice at 12–13 weeks of age as previously described [Tatsukawa *et al*., 2018; Yamagata *et al*., 2020]. Mice were placed on a perforated platform in a sealed Plexiglas chamber and heated by airflow from below. After a 5-min acclimation at room temperature, body temperature was increased by raising the air temperature at 0.5^°^C/min. Rectal temperature was continuously monitored, and baseline and seizure onset temperatures were recorded. Upon onset of a generalized seizure, mice were immediately transferred to an ice-cooled chamber until normothermia was restored. Seizure assessments were performed by an experimenter blinded to genotype and treatment.

## Acknowledgments

The authors thank the members of the Division of Genetic Therapeutics at Jichi Medical University, the Department of Neurodevelopmental Disorder Genetics, and the animal facility at Nagoya City University (NCU) for their support. We are grateful to the Research Equipment Sharing Center at NCU for technical assistance.

## Author Contributions

TY, TS and KY contributed to the study conception and design. Material preparation, data collection, and analysis were performed by TY, TS, HM, YH and KY. The first draft of the manuscript was written by TY, TS and KY. All authors have reviewed and approved the final manuscript.

## Data Availability

The datasets generated during and/or analyzed during the current study are available from the corresponding author on reasonable request.

## Declarations

### Ethics Approval

All animal breeding and experimental procedures were approved by the Institutional Animal Care and Use Committee of NCU (approval No. 19-032, approved 20 Dec 2022; approval No. 23-024, approved 21 Apr 2023). All procedures were conducted in accordance with the ARRIVE guidelines and the institutional guidelines and regulations of NCU.

### Consent to Participate

Not applicable.

### Consent for Publication

Not applicable.

